# RHOA mediated mechanical force generation through Dectin-1

**DOI:** 10.1101/663526

**Authors:** Rohan P. Choraghe, Tomasz Kołodziej, Alan Buser, Zenon Rajfur, Aaron K. Neumann

## Abstract

Dectin-1 is an innate immune pattern recognition receptor which recognizes β-glucan on the *Candida albicans* (*C. albicans*) cell wall. Recognition of β-glucan by immune cells leads to phagocytosis, oxidative burst, cytokine and chemokine production. We looked for specific mechanisms that coordinate phagocytosis downstream of Dectin-1 leading to actin reorganization and internalization of fungus. We found that stimulation of Dectin-1 by soluble β-glucan leads to mechanical force generation and areal contraction in Dectin-1 transfected HEK-293 cells and M1 macrophages. With inhibitor studies, we found this force generation is a SYK-independent, SFK (SRC Family Kinase)-dependent process mediated through the RHOA-ROCK-MLC pathway. We confirmed activation of RHOA downstream of Dectin-1 using G-LISA and stress fiber formation. Through phagocytosis assays, we found direct evidence for importance of RHOA-ROCK-MLC pathway in process of phagocytosis of *C. albicans*. In conclusion, we found evidence for RHOA-ROCK-MLC mediated mechanical force generation downstream of Dectin-1 for *C. albicans* phagocytosis.

## INTRODUCTION

Dectin-1 (Dendritic Cell associated C-type Lectin 1) is pattern recognition receptor expressed primarily on cells of myeloid origin including macrophages, dendritic cells and neutrophils^1^. The ligand for Dectin-1 is a highly immunogenic glucose polyscccharide with a backbone of β-(1–3)-linked linear units and β-(1–6) linked side chains known as β-glucan^2^. Glucans are ubiquitous in the fungal cell walls and form 50-60% of the dry weight of *Candida albicans* (*C. albicans*) cell walls^3^. Recognition of β-glucan by immune cells contributes to a variety of responses in these cells including non-opsonic phagocytosis, oxidative burst, regulation of transcription, production of inflammatory cytokines and chemokines, and initiation of adaptive immunity^4^. While Dectin-1 is expressed in up to eight different alternatively spliced isoforms, this paper will refer only to the predominant full length A isoform^5^. Downstream signaling elements activated upon recognition of β–glucan by Dectin-1 include SRC family kinases (SFK), SYK, and CARD9 to eventually mediate the production of reactive oxygen species, activation of NF-κB and subsequent secretion of pro-inflammatory cytokines^4^. Dectin-1 is known to signal via mechanisms that involve phosphorylation of the receptor’s cytoplasmic tail by SFK, leading to recruitment of SYK to the activated Dectin-1 cytoplasmic tail. SYK is then activated and phosphorylates downstream targets. Two major pathways are activated in a SYK-dependent manner: a pathway that leads to store operated calcium release, resulting in NFAT activation, and a PKC-dependent pathway that leads to activation of a CARD9 dependent signaling complex, which results in NF-kB activation^4,6,7^. Also, there is a SYK-independent pathway operated through RAS and RAF-1, leading to activation of MAPKs^8^.

Phagocytosis is a mechanism by which a cell engulfs particles typically >0.5 μm. In opsonic phagocytosis, a particle is coated by IgG and/or complement before being phagocytosed^9^. Opsonic phagocytosis by these two opsonins involves different patterns of RHO-family GTPase activation and actin reorganization to complete engulfment. Type-I opsonic phagocytosis by FcgR involves pseudopod extension and “zippering” of pseudopodial membrane extensions around particle. In contrast, type II opsonic phagocytosis through binding of complement receptors, such as integrin α_M_β_2_ (CR3, Mac-1, CD11b/CD18), to C3b and its proteolysis product invokes a phagocytic process characterized by “sinking” of the particle into cell body without obvious pseudopod extension^10^. These rapid and dramatic changes in local cellular morphology require actin reorganization driven by RHO-family small GTPases. During type I phagocytosis local pseudopod formation and membrane ruffling is thought to be mediated through RHO-GTPases such as RAC1 and CDC42. In contrast, sinking of particles in type-II phagocytosis is thought to be mediated by contraction of actomyosin network in RHOA dependent manner^10–12^.

Cells of the innate immune system can also engulf particles through non-opsonic phagocytic processes, such as recognition of β-(1,3)-glucan on *Candida* species fungal pathogens by Dectin-1^13,14^. Additionally, direct interaction of integrin α_M_β_2_ via its unique lectin-like domain is also important for *Candida* uptake by human neutrophils^15^, and signaling cross-talk between Dectin-1 and integrin α_M_β_2_ has been described^16^. But overall, Dectin-1 mediated phagocytosis is thought to be the major mechanism of non-opsonic recognition of glucan in macrophages and dendritic cells^13,17^.

Relative to opsonic phagocytic mechanisms, the specific signal transduction mechanisms leading to RHO-family GTPase activation and actin reorganization in Dectin-1 mediated phagocytosis have received less attention. Previous studies of Dectin-1 mediated phagocytosis of zymosan in murine macrophages have suggested that RAC2 and CDC42 are required, consistent with the activation of the Guanine nucleotide exchange factor (GEF) VAV1^14^. Previous studies have also found that non-opsonic recognition through Dectin-1 is SYK-independent^14,18^. However, SYK is required for RHO-GEF VAV1 activation during glucan phagocytosis in microglia^19^ and there is evidence for variable involvement of SYK in Dectin-1 mediated effector functions between various cell types^14,20^.

In this study, we present evidence that Dectin-1 activation by β-(1,3)-glucan causes SYK-independent activation of the small GTPase RHOA. The consequent actomyosin dependent contraction of the cellular cortex leads to substantial mechanical force generation downstream of Dectin-1. Dectin-1 and RHOA mediated generation of mechanical force through actomyosin networks is significant for non-opsonic engulfment of fungal particles and likely to contribute to the organization of the phagocytic synapse^21,22^.

## RESULTS

### Dectin-1 mediated, glucan-dependent cellular contraction

We observed that Dectin-1 expressing HEK-293 cells undergo areal contraction within minutes after stimulation with MMW (Medium molecular weight, ~150 kDa) glucan (Fig. 1a). MMW glucan is thought to mimic some structural characteristics of glucans in the cell wall, including a higher degree of triple helical structure relative to low molecular weight glucans such as laminarin. We found a statistically significant difference in control vs. Dectin-1 expressing cell area starting around 180 seconds after stimulating with MMW glucan (Fig. 1b). To further test our hypothesis in a system chosen for physiological relevance, we performed similar experiments in M1 macrophages differentiated from human peripheral blood monocytes^13,23,24^. Areal contraction similar to that observed with Dectin-1 expressing HEK-293 cells was also observed in M1 macrophages (Fig. 1c,1d), indicating the relevance and existence of a glucan sensitive mechanical force generation pathway downstream of Dectin-1 in human primary macrophages.

**Fig. 1.**
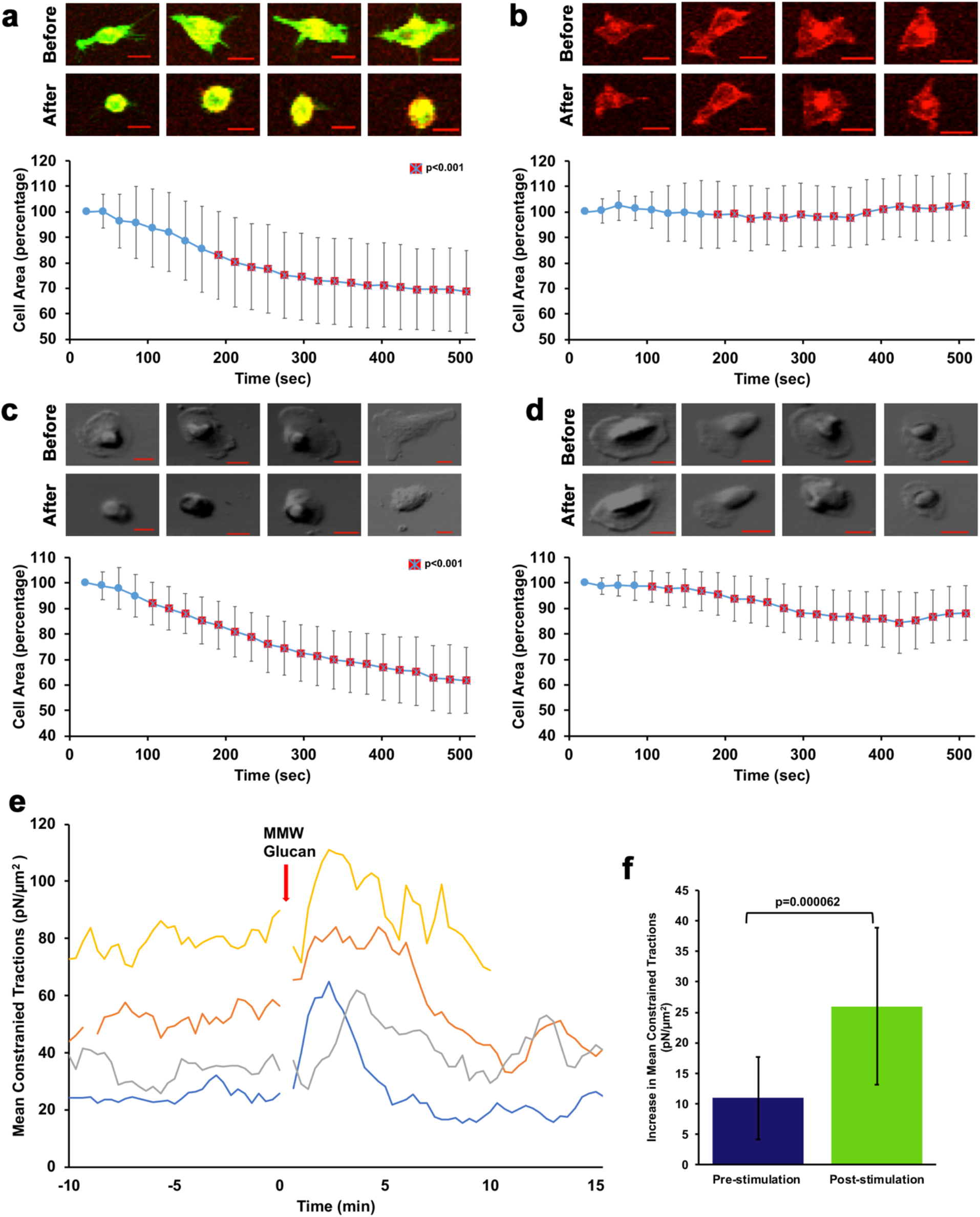
Dectin-1 mediated, glucan-dependent cellular contraction. **a** HEK-293 cells transfected with Emerald-Dectin-1 (green) and stained with Cell mask deep red (CMDR). Images labelled ‘Before’ are pre-stimulation and those labelled ‘After’ are 500 seconds after stimulation with MMW glucan. Scale bar, 20 μm. MMW glucan was added at time 0. (n=29) Data graphics in panel a-d represent sample mean and standard deviation. Red square with cross indicates significant difference in cell area at a timepoint between Dectin-1 transfected cells and control cells in panel 1b. **b** Control HEK-293 cells stained with CMDR, pre-stimulation and 500 seconds after stimulation with MMW glucan. Scale bar, 20 μm. (n=31). **c** DIC images of M1 macrophages before and after stimulation with MMW glucan. Scale bar, 50 μm. (n=31). **d** DIC images of mock stimulated M1 macrophages. Scale bar, 50 μm. (n=28). **e** Mean constrained tractions in 4 independent HEK-293 cells transfected with Emerald-Dectin-1 and stimulated with MMW glucan. **f** Average increase in mean constrained tractions of individual cells with standard deviation pre and post-stimulation with glucan (n=15).

Cells generate and apply mechanical forces to their environment, typically in the pN to nN range. Phagocytes exhibit an increase in cortical tension during the process of phagocytosis, exerting forces upon particles during their engulfment ^25^. One way to show and measure mechanical force application to the cellular microenvironment is traction force microscopy (TFM). With TFM we can measure these forces as local shear stresses (i.e., traction forces exerted by a cell on elastic substrate). We used TFM to measure the additional stresses exerted on the substrate by Dectin-1 expressing HEK-293 cells upon stimulation with MMW glucan. We found that 60% of cells showed increased tractions after stimulation with MMW glucan. Adherent cells undergo cycles of contraction and relaxation as they spread and migrate. Cells that did not generate measurable changes in tractions upon glucan treatment may have exhibited adhesion/migration-associated forces that masked glucan-triggered contraction. Figure 1e shows the TFM data of several representive cells exerting increased mean constrained tractions on the substrate after stimulation with MMW glucan. Increased mean constrained tractions after stimulation with MMW glucan over pre-stimulation tractions was found to be 15.02 pN/μm^2^ from 15 cells (Fig. 1f). These data confirm that significant mechanical forces are generated downstream of Dectin-1 activation by fungal glucan.

While mechanical force generation is clearly important for any phagocytic process, a specific signaling mechanism driving Dectin-1 dependent contraction of cortical actomyosin networks has not been clearly delineated. We hypothesized that such a signaling mechanism must exist in order to explain the observed Dectin-1 mediated contractile phenomenon. Integrin activation has been associated with RHOA activation and actomyosin contraction. However, HEK-293 cells do not express the glucan binding integrin α_M_β_2_^26^. Furthermore, control HEK-293 cells not expressing Dectin-1 failed to exhibit contraction in response to glucan stimulation (Fig.1b). Therefore, we conclude that Dectin-1 does trigger a cellular mechanical force generation response, and we focused on understanding Dectin-1 signaling mechanisms required to initiate this response.

### Biochemical signaling pathway downstream of Dectin-1 involved in force generation

We first tested the requirement for several membrane-proximal kinases in areal contraction mediated by MMW glucan ligation of Dectin-1 (Fig. 2a). The most membrane-proximal kinases in Dectin-1 pathway, SFK, were required to observe areal contraction after MMW glucan stimulation of Dectin-1 expressing cells. Treatment with PP1, SRC inhibitor almost completely abolished the contractile response. Treatment with a broad-spectrum PKC inhibitor had no impact on the Dectin-1 mediated areal contractility response, demonstrating that the PKC/CARD9 arm of Dectin-1 signal transduction is dispensable for mechanical force generation. Additionally, CARD9 is very poorly expressed in HEK-293 cells, suggesting that this pathway is minimally operative, if at all, in this model^27^. Interestingly, SYK inhibitor caused no significant decrease in Dectin-1 mediated cell contractility, suggesting that the membrane proximal signaling events leading to mechanical force generation downstream of Dectin-1 are SYK-independent. Because known SYK-independent signaling of Dectin-1 involves RAF-1 activation, which can lead to ERK MAPK activation, we inhibited the MAPKK MEK1&2 that activate ERK MAPKs. Disruption of this SYK-independent MAPK signaling pathway had no significant impact on cellular areal contraction. We concluded that mechanical force generation downstream of Dectin-1 requires a SFK-dependent process that does not appear to be mediated by any major known signaling mechanisms of Dectin-1.

**Fig. 2.**
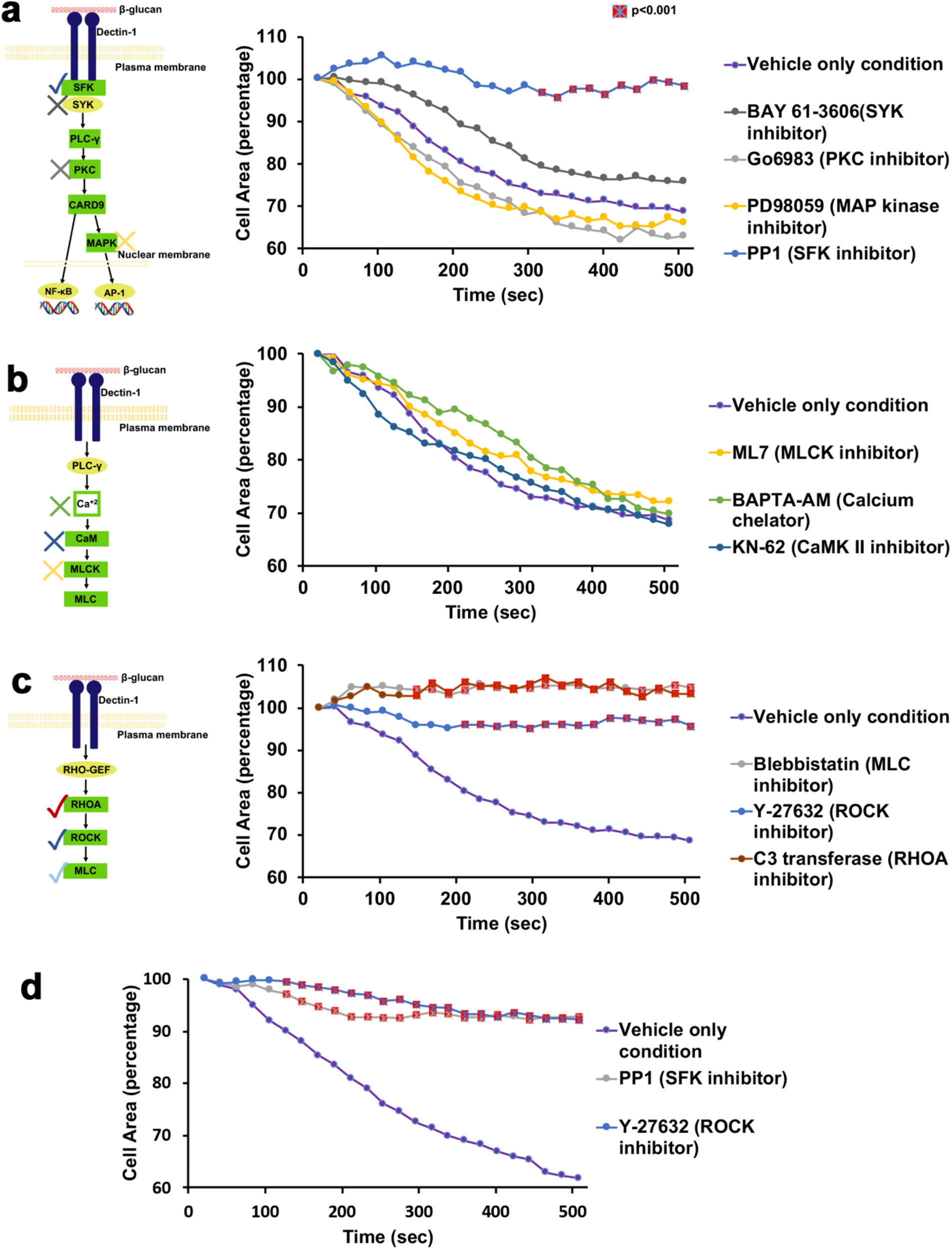
Biochemical signaling pathway downstream of Dectin-1 involved in force generation. **a** HEK-293 cells transfected with Dectin-1; pretreated with either BAY-61-3606 (SYK inhibitor) 500 nM (n=26), Go6983 (PKC inhibitor) 2 μM (n=24), PD98059 (MAP kinase inhibitor) 12.5 μM (n=26), PP1 (SRC inhibitor) 10 μM (n=28) for 1 hour; stimulated with 8 μg/ml MMW glucan and observed for change in area over 500 seconds. Red square with cross indicates significant difference in cell area at a time point between inhibitor condition and vehicle only condition. **b** HEK-293 cells transfected with Dectin-1; pretreated with either ML7 (MLCK inhibitor) 25 μM (n=36), BAPTA-AM (Calcium chelator) 5 μM (n=23), KN-62 (Ca^+2^/calmodulin-dependent kinase type II inhibitor) 20μM (n=23) for 1 hour; stimulated with 8 μg/ml MMW glucan and observed for change in area over 500 seconds. **c** HEK-293 cells transfected with Dectin-1; pretreated with either Blebbistatin (Myosin II inhibitor) 12.5 μM (n=35), Y-27632 (MLCK inhibitor) 5 μM (n=35) for 1hour or C3 transferase (RHOA inhibitor) 1.5 μg/ml (n=25) for 2 hours; stimulated with 8 μg/ml MMW glucan and observed for change in area over 500 seconds. **d** M1 macrophages pretreated with either PP1 (SRC inhibitor) 10 μM (n=24), Y27632 (ROCK inhibitor) 5 μM (n=25) for 1 hour; stimulated with 8 μg/ml MMW glucan and observed for change in area over 500 seconds.

Because no known Dectin-1 intracellular signaling pathways are clearly connected to cellular force generation mechanisms that could explain the observed contractile phenomenon, we proceeded to broadly consider cellular mechanisms of mechanical force generation. Cellular force generation proceeds through two main effector mechanisms, both of which finally impinge on actin-myosin based contraction^12^. First, muscle cells generate force via a calcium/calmodulin dependent process that leads to Myosin Light Chain Kinase (MLCK) activation, phosphorylation of myosin light chain (MLC) and activation of actomyosin contraction. Second, other cell types utilize a different pathway (e.g., for cell migration) that involves activation of RHOA, leading to RHO-associated protein kinase (ROCK) activation, phosphorylation of myosin light chain and initiation of actomyosin contraction. We found that inhibition of the calcium/calmodulin pathway using several inhibitors against distinct steps in this pathway uniformly showed no significant effect on the Dectin-1 mediated contractile response (Fig. 2b). This led us to conclude that the first pathway (calcium/calmodulin dependent) was not involved in the Dectin-1 mediated pathway leading to force generation. Next, we tested inhibitors against RHOA and ROCK to determine the involvement of the second pathway. We found that these inhibitors, as well as inhibitor for myosin II itself, were strongly inhibitory for the Dectin-1 mediated contractile response (Fig. 2c). We concluded that the RHOA based force generation pathway was engaged downstream of Dectin-1 activation by glucan, leading to myosin II activation for actomyosin contractility. Further we also confirmed existence this force generation pathway downstream of Dectin-1 in M1 macrophages. SRC inhibitor and ROCK inhibitor strongly inhibited contractility in macrophages (Fig. 2d) indicating the importance of these pathways in a physiologically relevant system.

### Activation of RHOA downstream of Dectin-1 after stimulation with glucan

Activation of small GTPases, such as RHOA, RAC1 and CDC42, gives rise to typical morphological responses, including formation of actin stress fibers, lamellipodial/pseudopodial extension or filopodial extension, respectively^28^. Dectin-1 transfected cells, on stimulation with glucan, predominately showed increased formation of actin stress fibers, a phenotype consistent with activation of RHOA (Fig. 3a). Supporting this conclusion, we observed an increase in active RHOA within minutes of Dectin-1 stimulation by G-LISA assay (Fig. 3b). These observations confirm and support our conclusion that Dectin-1 stimulation by soluble MMW glucan does reliably induce a contractile response of the cell body, and that this force generative response depends upon SFKs, the small GTPase RHOA, ROCK and myosin II (Fig. 3c).

**Fig. 3.**
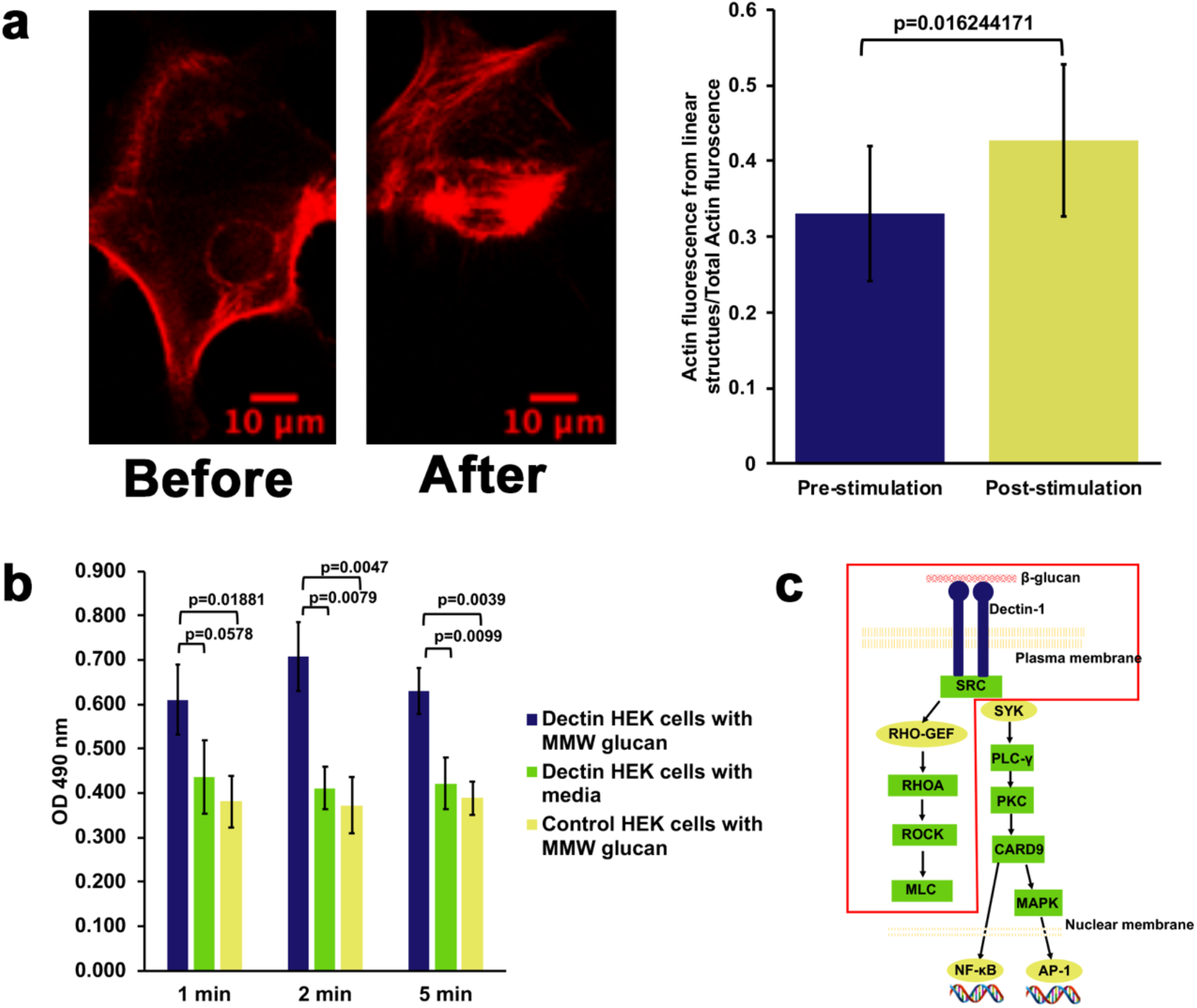
Activation of RHOA downstream of Dectin-1 after stimulation with glucan. **a** HEK-293 cell transfected with Emerald-Dectin-1 and mCardinal-LifeAct showing increased actin stress fiber formation 5 minutes after stimulation with MMW glucan. Graph represents average fraction of actin in linear form with standard deviation before and 5 minutes after stimulation with MMW glucan. (n=13). **b** RHOA G-LISA showing significant activation of RHOA after stimulation of Dectin-1 transfected HEK-293 cells with MMW glucan compared to mock stimulation or Control HEK-293 cells stimulated with MMW glucan. Bars represent average OD with standard deviation for each condition for 3 independent experimental replicates. **c** Final schematic of force generation pathway downstream of Dectin-1.

### Relevance of force generation pathway in non-opsonic phagocytosis mediated by Dectin-1

Next, we examined the importance of mechanical force generation in non-opsonic phagocytosis downstream of Dectin-1. HEK-293 cells transfected with Dectin-1 were presented with a high glucan-exposure *C. albicans* clinical isolate strain, TRL035, stained with calcofluor and Cypher5E, a dye which registers entry into the acidic phagolysosome via a substantial increase in fluorescence. Following 1 hour of exposure to yeast in the presence or absence of inhibitors of the RHOA pathway, both the phagocytic index (Fig. 4a), a measure of the efficiency of phagocytosis for captured fungal particles, and phagocytic activity, the fraction of cells that phagocytosed yeast were calculated. Inhibition of RHOA, ROCK, or myosin II significantly decreased both measures of Dectin-1 dependent phagocytosis, confirming an important role for this contractility signaling pathway in Dectin-1 mediated phagocytosis. Using the human M1 macrophage system, we performed similar phagocytosis assay with TRL035 *C. albicans* and demonstrated a significant decrease in phagocytosis with RHOA inhibitor, indicating importance of the RHOA based pathway for phagocytosis of fungi in macrophages (Fig. 4b).

**Fig. 4.**
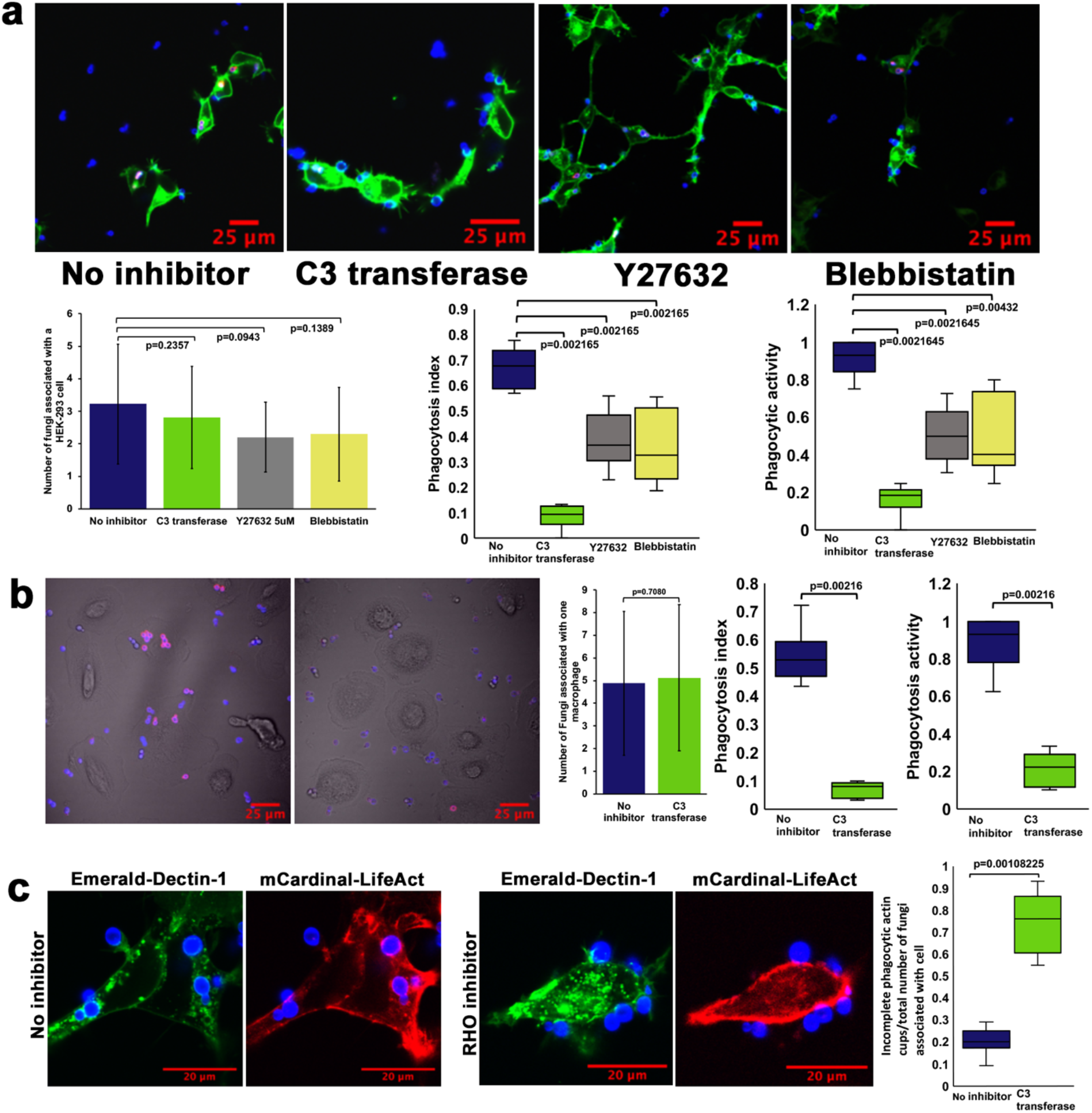
Relevance of force generation pathway in non-opsonic phagocytosis mediated by Dectin-1. **a** Phagocytosis assay with HEK-293 cells transfected with Emerald-Dectin-1 and presented with TRL035 *Candida albicans. C. albicans* were stained with Calcofluor white (Blue) and Cypher5E (Red). Images were taken 1 hour after introduction of *C. albicans* to culture dishes. HEK-293 cells were either under no inhibition condition (n=68) or pretreated with C3 transferase 1.5 μg/ml (n=38) for 2 hours, Blebbistain 12.5 μM (n=43) or Y27632 5 μM (n=69) for 1 hour. Graphs in panel a and b show the average number of fungi associated with a single cell (mean±S.D.), Phagocytosis index and Phagocytic activity under each condition. **b** Phagocytosis assay in M1 macrophages with TRL035 *C. albicans* under no inhibitor (n=52) and C3 transfearse 1.5 μg/ml (n=59) conditions. **C** Phagocytosis assay in HEK-293 cells transfected with Emerald-Dectin-1 and mCardinal-LifeAct with TRL035 *C. albicans*. Number of fungi associated with incomplete phagocytic actin cups compared to total number of fungi associated with cell, under no inhibitor (n=44) and C3 transferase 1.5 μg/ml (n=49) condition were counted and are shown in the graph.

We observed that, while Dectin-1 mediated phagocytosis is markedly reduced in cells subjected to inhibition of the ROCK/RHOA pathway, partial actin phagocytic cups were observed in 75% of contacts with yeast in RHOA inhibited cells (Fig. 4c). This suggested the RHOA activity might be required for complete maturation of the phagocytic synapse, leading to closure of the cup, particle engulfment and formation of the phagolysosome.

## DISCUSSION

Dectin-1 engages both SYK-dependent and -independent signaling pathways, and prior literature suggests some context-dependent variability in the signaling mechanisms engaged by this important anti-fungal receptor. Dectin-1 mediated activation of NF-κB in a SYK dependent manner, leading to formation of reactive oxygen species and other immune responses, is very well documented^4^. In contrast Gringhuis et al.^8^ showed that Dectin-1 also induces a SYK-independent activation of NF-κB through RAF-1 to control adaptive immune responses. In similar fashion, as mentioned by Goodridge et al.^18^, literature about signaling downstream of Dectin-1 giving rise to phagocytosis is also variable. Even though Dectin-1 signaling clearly leads to activation of SYK, it does not appear to be important in phagocytosis as indicated by our data. Herre et al.^14^ similarly found that phagocytosis downstream of Dectin-1 is SYK-independent. We also found that mechanical force generation required for phagocytosis downstream of Dectin-1 is SYK-independent. We demonstrated that Dectin-1 dependent force generation pathway of phagocytosis is mediated through SFK-dependent activation of yet unknown RHO-GEF leading to activation of RHOA and ROCK, ultimately leading to the formation and contraction of an actomyosin network. RHOA mediated contractility during Dectin-1 activation allows the application of forces on the order of 15 pN/μm^2^ by the cell on its environment.

We found RHOA plays a very important role in Dectin-1 mediated force generation through actomyosin network formation and contraction in phagocytosis of *C. albicans*. In contrast, Herre et al.^14^ found that CDC42 and RAC1 but not RHOA play significant role in phagocytosis downstream of Dectin-1. Their system was different than ours as they used murine NIH/3T3 fibroblast cell line with zymosan particles whereas we used human cell line models and primary human macrophages with a *C. albicans* clinical isolate strain. Therefore, this discrepancy may be explained by Herre et al’s use of murine cell background and use of particles with much higher level of glucan exposure.

RHO-family GTPase usage varies widely between phagocytic contexts and mechanisms, but our data suggest that a type II-like process in engaged by Dectin-1. In phagocytosis of apoptotic cells, Tosello-Trampont et.al^29^ and Nakaya et al.^30^ found that RHOA and ROCK have an inhibitory effect on phagocytosis. They found that CDC42 plays a role in phagocytosis of apoptotic cells, and this suggests that CDC42 based type-I phagocytosis might be involved in phagocytosis of apoptotic cells compared to involvement of RHOA and ROCK in type II phagocytosis^12^. In Dectin-1 based phagocytosis of *C. albicans*, we think type II sinking phagocytosis is predominantly involved. In sinking phagocytosis, RHOA plays important role in formation of actomyosin network formation. In a mechanistically parallel system, Colucci-Guyon et al.^31^ and Wiedemann et al.^32^ showed that localized actin assembly which acts as the driving force for particle engulfment in CR3-mediated type-II phagocytosis is mediated by RHOA. Further studies will continue to clarify differential use of phagocytic mechanisms and dependence upon various small GTPases for variety of relevant cell types, particles and phagocytic routes.

One explanation for why different phagocytosis machinery is used in type-I versus type-II phagocytosis could be differences in final desired result of phagocytosis. Erwig et al. ^33^ found that macrophages engulfing apoptotic cells engaged a rapid, RHOA-dependent process of phagosomal acidification that was much faster than the rate of phagosome acidification in macrophages engulfing Ig-opsonized particles or DCs engulfing either type of particle. The authors proposed a differential role of RHOA in phagosome maturation depending upon whether the cell type involved was specialized for degradation/eradication versus antigen acquisition from the ingested particles.

Kim et al.^34^ reported that ROCK inhibition increased phagocytosis of apoptotic cells suggesting an initial inhibitory effect of RHOA on phagocytosis in this system. However, this was later followed by transient RHOA activation prior to closure of phagocytic cup. Transient RHOA activation prior to closure of phagocytic cup was hypothesized as an important regulatory step in the “don’t eat me” signaling pathway. Our findings suggest that RHOA may be similarly important for phagocytic cup closure in the Dectin-1 system. Future studies will be necessary to determine whether RHOA activation is also important for earlier stages of Dectin-1 dependent engulfment of fungal pathogens. In terms of mechanics of phagocytosis as mentioned by Herant et al.^25^, the final phase of phagocytosis should involve a “pull-in” force, probably mediated by myosin II. In our study, we found Dectin-1 based phagocytosis of *C. albicans* involves mechanical force generation through RHOA and myosin II. This mechanical force should provide the required pull in force for phagocytosis in the final phase of Dectin-1 mediated engulfment.

One more factor which could be driving differential actomyosin machinery in phagocytosis could be particle size. As mentioned by Clarke et al.^35^ and Schlam et al.^36^ size of the engulfed particle could play role in actomyosin responses. In case of large particles (>10 um), protrusive responses could play more important role whereas in smaller particles (<5 um), pulling in force could play more important role. Overall it appears that RHO family small GTPases play differential roles in the process of phagocytosis depending on type of phagocytosis, size of particle being phagocytosed and ultimate desired result of phagocytosis. Such findings might predict a stronger role for RHOA in phagocytosis of *C. albicans* yeast-sized fungi, but perhaps a greater requirement for other RHO-family GTPases during internalization of hyphae or larger fungal cells/structures.

Finally, the actomyosin network formation is very well recognized to be involved in the movement of T cell receptor (TCR) microclusters and immunological synapse formation^21^. We propose that the actomyosin network could play important role in recruitment of innate immune receptors to the fungal contact sites. This model of fungal contact site formation comprises several steps: initial pathogen capture, early Dectin-1 activation, generation of a RHOA dependent actomyosin flow at the contact site, and recruitment of additional Dectin-1 for amplified and prolonged signaling leading to phagocytosis. Initial pathogen capture is likely dominated by interactions between abundant cell wall surface mannans and immunoreceptors with relevant binding specificities (e.g., DC-SIGN, CD206). However, Dectin-1 mediated phagocytosis requires this receptor to find its much more sparsely exposed ligand on the cell wall (i.e., nanoscale glucan exposures)^37^. We propose that sparse Dectin-1/glucan interactions may be rate limiting for phagocytosis in physiologically relevant settings. So, initial detection of glucan by Dectin-1 initiates an actomyosin flow that recruits additional Dectin-1 to the contact site, maximizing the probability that available glucan exposure sites will be recognized so that phagocytic response can be triggered. This actomyosin flow may also recruit more mannan-binding receptors to provide better particle retention by the phagocyte. Future studies will be required to test the role of actomyosin flow at the pathogen contact site in organization of the phagocytic synapse and initiation/amplification of signaling leading to fungal pathogen phagocytosis.

### Methods

#### Cell culture

HEK-293 cells (ATCC, #CRL-1573) were cultured in DMEM, 10% FBS, 1% Penicillin-Streptomycin, 2mM L-glutamine, 1mM Sodium pyruvate medium at 37°, 5% CO_2_ environment in an incubator.

#### Transfection

Emerald-Dectin1A-N-10 was a gift from Michael Davidson (Addgene plasmid, #56291; http://n2t.net/addgene:56291; RRID: Addgene_56291). Transfection with Emerald-Dectin-1A-N-10 was performed using standard protocols of Fugene 6 (Promega, #E2691). Cells were selected using Geneticin (G418 Sulfate) (Thermo-Fischer 10131035) at 400 μg/ml for 2 weeks.

#### M1 macrophages

De-identified Primary Peripheral Blood Mononuclear Cells (PBMC) (ATCC, #PCS-800-011) were thawed and cultured using Roswell Park Memorial Institute (RPMI) 1640 medium, 10% FBS, 1% Penicillin-Streptomycin, 2 mM L-glutamine, 1 mM Sodium pyruvate medium at 37°, 5% CO_2_ environment in incubator^38^. For generation of M1 macrophages, 50 ng/mL rhGM-CSF (PeproTech, #300-03) was added to culture dish for 7 days followed by stimulation with 500 ng/mL LPS (Invivogen, #tlrl-peklps) for 24 hours^24,38^.

#### MMW glucan

β-1,3;1-6-glucan from *S. cervisiae* cell wall extraction (~150 kDa) was a generous gift from ImmunoResearch Inc. (Eagan, MN). Glucan was weighed and resuspended in respective cell culture media of cell line to be stimulated.

#### Contractility assay

HEK-293 cells or M1 macrophages were grown in 35 mm glass bottom Mattek dishes. Cells were stained with CellMask Deep Red Plasma Membrane stain (CMDR) (Invitrogen, #C10046) (1:100 dilution in HEK cell media) at 10 μl/ml of media in dish for 1 hour. Cells were then stimulated with MMW glucan at 8 μg/ml and imaged in FV1000 laser scanning confocal microscope (Olympus, Center Valley, PA) with controlled temperature, 37°, 5% CO_2_ for 10 minutes. 10x lens, 0.40 NA, Plan-Apochromat objective lens was used for imaging. Emerald-Dectin-1 was excited with a 20 mW, 473 nm diode laser operated at 1% power and CMDR was excited with a 20 mW, 635 nm diode laser operated at 2% power. These lines were reflected to the specimen by a 405/473/559/635 multi-edge main dichroic element, routed through confocal pinhole (110 μm diameter) to dichroics followed by bandpass emission filters in front of 2 independent High sensitivity GaAsP PMT detector (HSD). Specifically, the emission light passed by the main dichroic was directed to HSD1 (Emerald-Dectin-1 channel) via SDM560 filter cube and passage through a BA490-540 nm bandpass filter. For CMDR channel from SDM560 filter cube to BA575-675 nm bandpass filter. Different conditions used were either no inhibitor condition or pretreatment with various inhibitors for 1 hour; BAY-61-3606 (EMD Millipore, #574714) 500 nM^39^, Go6983 (Cayman Chemical Company, #13311) 2 μM^40^, PD98059 (Invivogen, #tlrl-pd98) 12.5 μM^41^, PP1 (Cayman Chemical, # 4244) 10 μM^42^, ML7 (Sigma-Aldrcih, #I2764) 25 μM^43^, BAPTA-AM (Cayman Chemical, #15551) 5 μM^44^, KN-62 (Cayman Chemical, #13318) 20 μM^45^, Blebbistatin (Sigma-Aldrich, #203390) 12.5 μM, Y-27632 (Sigma-Aldrich, #Y0503) 5 μM^46^. For RHO inhibitor condition cells were treated with C3 Transferase (Cytoskeleton Inc., #CT04) 1.5 μg/ml for 2 hours.

#### TFM

TFM gels were prepared similarly to Valon et al.^47^. Briefly, the polymerization mixture was placed in glass-bottom dish and covered with 18 mm round coverslip to polymerize for 1 hour. Then, the substrates were kept overnight in PBS in 4°C to wash out any potentially non-polymerized acrylamide monomers from the substrate. Next day the substrates were activated using Sulfo-SANPAH, covered with extracellular matrix (ECM) proteins and incubated for 20 hours in 4°C. Afterwards, dishes with substrates were washed 3 times with PBS and cells were seeded on substrates.

The first step of Traction Force Microscopy experiments was substrate optimization. To find optimal elasticity and suitable ECM protein cover, substrates of 3 different elasticities (1 kPa, 8.4 kPa, 19 kPa), similarly to ones defined in Tse & Engler et al.^48^, and two ECM proteins (Fibronectin & Type I Collagen) were tested. The aim of this optimization was to find substrates which are soft enough to observe significant cell tractions, but stiff enough to keep cells spread on the substrate. Cells were observed after 3, 6 and 24 hours after seeding. After 3 and 6 hours, cells were spread on all mentioned substrates, however after 24 hours cells plated on fibronectin-coated substrates were mostly detached from the substrate, while cells seeded on collagen coated substrates were still attached to the substrate. Cells grown on collagen-coated substrates of elasticities 19 and 8.4 kPa were well spread, while those on soft substrate (1 kPa) were poorly spread, which mean they were not strongly attached to the substrate. That fact suggested that we would not be able to observe significant traction forces after stimulation. This optimization led us to the conclusion that optimal substrate to be used in our TFM experiments was 8.4 kPa coated with type-1 collagen.

TFM experiments were performed 18 hours after cell seeding, using Zeiss Axio Observer Z1 inverted epifluorescence microscope. In each experiment, 4 fields of view were observed every 20 seconds. The aperture of fluorescent light was constrained only to the observed field of view so as to minimize possible cytotoxic effects while illuminating the whole sample with fluorescent light for longer time. The continuous TFM experiments were performed for 25 minutes with an interval of 20 seconds. 10 minutes after start, the solution of MMW glucan was added to reach the final concentration of 8 μg/ml in cell medium and the cells responses were observed for next 15 minutes. Analysis of TFM experiments was performed using software provided by X.Trepat, Institute for Bioengineering of Catalonia, Barcelona, Spain^47^.

For calculating increase in tractions following formulae were used

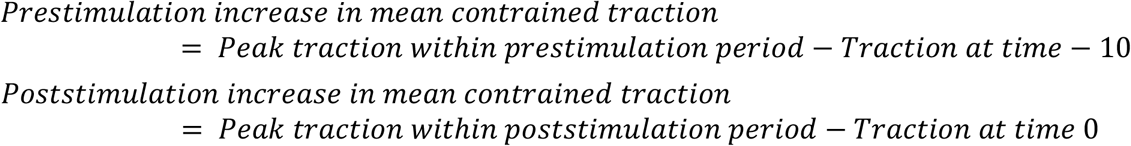

For calculating fraction of cells which showed increase in traction after stimulation with MMW glucan (60% in our experiments), cells which showed 50% or more increase in tractions over pre-stimulation increase were considered as cells showing positive response. Only cells showing positive response were considered for calculating actual value (15.02 pN/μm^2^ in our experiments) for increase in traction after stimulation with glucan.

#### Actin-stress fiber imaging

HEK-293 cells stably expressing Emerald-Dectin-1 were cotransfected with mCardinal-LifeAct-7 using Fugene 6 (Promega#E2691) protocol. mCardinal-Lifeact-7 was a gift from Michael Davidson (Addgene plasmid # 54663; http://n2t.net/addgene:54663; RRID:Addgene_54663). Cells were stimulated with glucan at 8 μg/ml and imaged in Olympus FV1000 confocal system (optical description similar to contractility assay except for using 60x super-corrected, 1.40 NA, Plan-Apochromat oil immersion objective) with controlled temperature, 37°, 5% CO_2_ for 10 minutes. Actin stress fibers were quantified using FSegment matlab script developed by Rogge et al.^49^ Fraction of actin in linear form was quantified before stimulation and 5 minutes after stimulation.

#### RHOA G-LISA

G-LISA RHOA activation assay kit (Cat. #BK124) from Cytoskeleton, Inc. was used. Dectin-1 transfected HEK-293 cells stimulated with 8 μg/ml MMW glucan were analyzed for RHOA activation at 1 minute, 2 minutes and 5 minutes. Control conditions were Dectin-1 transfected cells stimulated with media (vehicle) and Control HEK-293 cells stimulated with MMW glucan.

#### Fungal culture

Clinical isolate of TRL035 *C. albicans* was obtained as previously decribed^50^. Isolate was stored as glycerol stock in −80°C freezer. This stock was transferred to 5 ml filter-sterilized yeast extract-peptone-dextrose (YPD) medium (Becton Dickinson) and grown for 16 h at 30°C, with a shaking speed of 300 rpm. Such growth conditions yielded yeast cells at the late log phase^51^.

#### Phagocytosis assay

TRL035 *C. albicans* were first stained with Fluorescent Brightener 28 (Calcofluor white) (Sigma-Aldrich #F3543). 25 μl of 1 mg/ml calcofluor white was used to stain 1 ml of 16-hour fungal culture in PBS (Gibco) pH 7.4 for 15 minutes. Then cells washed 3 times with PBS and stained with 75 μM CypHer5E NHS ester (GE Healthcare, PA #15401) for 1 hour at 25°C^50^. *C. albicans* were then added to culture dishes and observed under microscope for increased fluorescence in CypHer5E channel to look for fungi which had been phagocytosed. Images were taken with a FV1000 laser scanning confocal microscope (Olympus, Center Valley, PA) equipped with a 60x super-corrected, 1.40 NA, Plan-Apochromat oil immersion objective. Calcofluor White (marker for all yeast) was excited with a 50 mW, 405 nm diode laser operated at 1% power and Cypher5E was excited with a 20 mW, 635 nm diode laser operated at 0.5% power. These lines were reflected to the specimen by a 405/473/559/635 multi-edge main dichroic element and routed through confocal pinhole (110 mm diameter) to secondary dichroics followed by bandpass emission filters in front of 2 independent PMT detectors. Specifically, the emission light passed by the main dichroic was directed to PMT1 (Calcafluor white channel) via reflection from the SDM473 dichroic and passage through a BA430-455 nm bandpass filter. For Cypher5E channel (HSD detector) light from SDM473 was directed to SDM560 filter cube to BA575-675 nm bandpass filter. Phagocytosis index and Phagocytic activity were calculated using formulae below. Various inhibitor conditions used were same as in contractility assay.

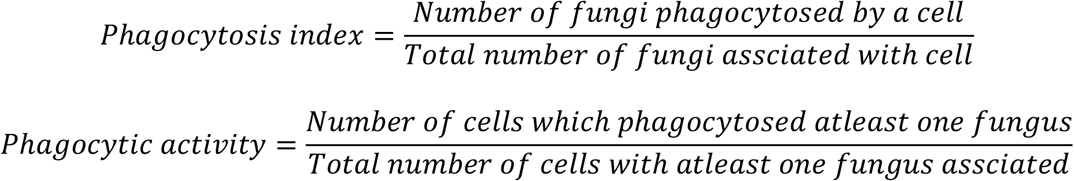

#### Statistical analysis

Statistical comparisons were carried out using an unpaired twotailed t-test for independent samples. Standard deviation for each sample and p-value for comparisons are indicated in figures. For phagocytosis index, phagocytosis activity and actin phagocytic cup index comparison, Mann-Whitney U test was used. Statistical analyses and graphing were performed by Microsoft Excel 2019 version 16.23.

## Acknowledgements

We gratefully acknowledge technical advice and critical reading of the manuscript by Akram Etemadi Amin, Eduardo Anaya and Carmen Martinez. Glucans used in this study were a generous gift of ImmunoResearch Inc, MN., which did not control experimental design, data interpretation or decision to publish. We thank Xavier Trepat and Carlos-Perez Gonzalez for providing training and technical advice with regard to TFM methodologies used in this work. The authors declare that they have no conflicts of interest relevant to this work. This research was supported by the University of New Mexico Center for Spatiotemporal Modeling of Cell Signaling (STMC; NIH P50GM085273, AKN) and R01AI116894 (AKN).

